# Versatile Modular Antibodies for Sensitive and Specific Detection of Poly-ADP-Ribose

**DOI:** 10.1101/2025.02.07.637082

**Authors:** Helen Dauben, Ivan Matić

**Affiliations:** Research Group of Proteomics and ADP-Ribosylation Signaling, Max Planck Institute for Biology of Ageing, 50931 Cologne, Germany; Cologne Excellence Cluster for Stress Responses in Ageing-Associated Diseases (CECAD), University of Cologne, 50931 Cologne, Germany

## Abstract

ADP-ribosylation is a chemically versatile modification of proteins, RNA and DNA that regulates various important signaling pathways, many of which are implicated in human diseases. Despite being discovered 60 years ago in the form of poly-ADP-ribosylation - the most studied form of this modification - investigating ADP-ribosylation at the molecular level has historically been challenging. By applying serine ADP-ribosylation-based antibody engineering technology, we have developed the first site-specific, as well as sensitive, mono-ADP-ribosylation modular antibodies. Here, we extend the scope of this technology to poly-ADP-ribosylation. By combining serine poly-ADP-ribosylated peptides as antigens, PARP1 serine mono- and poly-ADP-ribosylation for validation, phage display and the SpyTag protein ligation systems, we developed modular antibodies that are highly specific for poly-ADP-ribosylation. SpyTag-based coupling of horseradish peroxidase at specific positions, distant from the antigen-binding region of the antibody, yields a format that simplifies immunoblotting while dramatically enhancing poly-ADP-ribosylation detection sensitivity. Additionally, the creation of synthetic immunoglobulin formats – mouse, rabbit and human – enables straightforward co-detection of mono- and poly-ADP-ribosylation in cells by immunofluorescence. Our new tools improve and simplify the detection of poly-ADP-ribosylation, particularly in immunoblotting, enabling specific investigations of this key cellular signal in the context of other distinct forms of ADP-ribosylation.

## Introduction

ADP-ribosylation is a versatile post-translational modification (PTM) that regulates numerous cellular processes, including DNA repair, transcription, signal transduction, and cell death^1–4^. It involves the enzymatic transfer of ADP-ribose units from nicotinamide adenine dinucleotide (NAD⁺) to target molecules, mediated primarily by members of the ADP-ribosyltransferase (ART) enzyme family^5^. This dynamic modification can occur as mono-ADP-ribosylation (mono-ADPr), where a single ADP-ribose unit is present, or poly-ADP-ribosylation (poly-ADPr), involving the formation of linear or branched chains of ADP-ribose. The reversibility of this process, mediated by ADP-ribosyl hydrolases, underscores its regulatory significance in maintaining cellular homeostasis^6,7^. First identified in the 1960s, ADP-ribosylation has since emerged as a critical player in cellular stress responses, particularly in the context of DNA damage repair^8,9^. Poly(ADP-ribose) polymerase 1 (PARP1), the most studied member of the PARP family, is activated in response to DNA strand breaks, orchestrating the recruitment of DNA repair machinery and chromatin relaxation. Beyond DNA repair, ADP-ribosylation influences chromatin remodeling, RNA processing, and immune signaling. Notably, dysregulation of ADP-ribosylation is implicated in various diseases, including cancer, neurodegenerative disorders, and inflammatory conditions, making it an attractive target for therapeutic intervention^10–14^.

The biological complexity of ADP-ribosylation stems from the diversity of its enzymes, substrates, and cellular contexts. The PARP family alone consists of 17 members in humans, of which four are known to synthesize poly-ADPr (PARP1, PARP2, PARP5a/TNKS1, and PARP5b/TNKS2), each exhibiting distinct substrate specificities and cellular functions^15^. Moreover, emerging evidence highlights non-canonical roles of ADP-ribosylation, such as its involvement in liquid-liquid phase separation and the regulation of stress granules^16–18^.

ADPr is one of the most chemically complex PTMs, with conjugation chemistries ranging from O-glycosidic (serine and tyrosine), S-glycosidic (cysteine), N-glycosidic (arginine) to ester (aspartate and glutamate) linkages. Strikingly, some of the main conjugation chemistries have remained elusive for decades. A prime example of this is serine ADP-ribosylation, which, despite being the most abundant - though not the exclusive target residue in PARP1/2 signaling - was only discovered less than ten years ago^19–22^.

The structural complexity of ADP-ribosylation arises from the unique structure of ADP-ribose and its ability to form both mono-ADPr and intricate polymer chains. The polymeric forms, particularly poly-ADPr, can adopt linear or branched conformations, creating a diverse array of structural motifs^23,24^. These variations influence the binding affinity and functionality of effector proteins, known as “readers,” which recognize specific poly-ADPr structures through macrodomains, PAR-binding zinc fingers, and other motifs^25–28^. Additionally, the rapid synthesis and degradation of these chains add a temporal layer of complexity, making ADP-ribosylation a highly dynamic and versatile regulatory mechanism.

The inherent biological and chemical/structural complexity of ADPr has historically hindered its study and the development of appropriate analytical tools. This lack of tools has historically remained a significant limitation in many studies. Nevertheless, the critical role of poly-ADPr in modulating essential biological processes, particularly PARP1 signaling in the DNA damage response, has driven the development of various approaches and tools to study this modification since its discovery in the 1960s^29^.

ADPr research methods and tools can be broadly categorized into indirect and direct approaches. Indirect methods include the use of small molecule inhibitors to block poly-ADPr writers^30^, or genetic and RNA interference-based techniques to modulate the occurrence and persistence of this modification by targeting readers or erasers. Direct approaches aim to study poly-ADPr either *in-vitro* or *in-vivo*, including artificial construction or modification of poly-ADPr^31,32^ and its targets to unravel its structural complexity and biological functions. Early techniques utilized NAD⁺ analog probes^33^, which remain in use today with further optimizations, such as Et-DTB-NAD⁺^34–36^. In addition to chemical probes, enrichment strategies originally designed for other post-translational modifications (e.g., phosphoenrichment) have been adapted for poly-ADPr^37^. Elegant methods such as Enzymatic Labeling of Terminal ADP-Ribose (ELTA) offer another avenue for studying this modification by leveraging enzymes like OAS1 and modified dATP analogs to chemically tag and track poly-ADPr chains^38^.

Among direct methods to study endogenous poly-ADPr levels, antibodies and reagents specific to poly-ADPr have emerged as promising tools to advance the field. Notably, many of the most frequently cited poly-ADPr antibodies have been polyclonal, which are challenging to produce consistently, leading to their discontinuation. This left the field reliant on monoclonal antibodies, such as 10H and the much less commonly used 16B. The 10H antibody, which was the most widely used tool in the early decades of ADPr research, primarily recognizes long poly-ADPr chains starting at 10–20 units and has low sensitivity compared to reagents developed in the last decade.

Recently, the Kraus laboratory pioneered the conversion of established ADP-ribose reader domains into Fc fusion, antibody-like recombinant reagents. One widely used tool is the AF1521 domain, which has been applied in various studies. Its mutated from, with increased affinity is used particularly for enrichment followed by mass spectrometry analysis^39,40^. While this approach has identified numerous ADPr sites, it cannot distinguish between mono- and poly-ADPr because the domain binds to the terminal unit, enriching both forms indiscriminately. Under certain conditions, both the original and affinity-matured AF1521 domain exhibit hydrolytic activity towards aspartate and glutamate ADPr^39^. Another approach employs the WWE domain of RNF146, which specifically binds poly-ADPr. More precisely, it targets iso-ADPr, a structure unique to poly-ADPr chains, thereby avoiding misinterpretation by not interacting with mono-ADPr^41^. This innovation added a more selective tool for studying poly-ADPr^42,43^. However, the diversity of poly-ADPr-specific domains, each potentially evolved to recognize distinct poly-ADPr structures, suggests that individual domains might exhibit specificity for certain chain types or protein targets. Even among WWE domains, differences in binding specificity and affinity have been observed, as exemplified by the two WWE domains in PARP13^44^. As a result, the application of these protein domain-based tools may be beneficial to study specific research questions rather than offering the broad applicability of antibodies. In fact, a so-called ‘Poly/Mono-ADP Ribose’ rabbit monoclonal antibody is frequently used to detect poly-ADPr. While it is sensitive and specific for ADP-ribose, this antibody is not specific for poly-ADP-ribose, as it also recognizes mono-ADP-ribose^45^.

By converting our biological discovery of serine ADPr by HPF1/PARP1^19,20^ into a technology for generating ADPr peptides we have developed several anti-ADPr modular antibodies^21,46–49^. This includes the first - and so far only - site-specific ADPr antibodies^46^, as well as a sensitive and specific mono-ADPr antibody that has proven effective in detecting not only serine mono-ADPr^47^ but also beyond PARP1 signaling, including interferon signaling^50–52^. Importantly, serine ADPr reactions not only yield antigens for the initial selection of antibody candidates via phage display but also play a crucial role in subsequent steps, enabling in-depth validation and characterization of the best antibody candidates. While phage display used in our approach offers several advantages compared to traditional animal-based approaches for generating antibodies, including the rapid initial identification of multiple binders to the antigen of choice, it has the drawback of yielding recombinant antibodies in non-standard formats. While conversion to standard immunoglobulins and other formats is feasible, cloning-based conversion is impractical and time-consuming, counterbalancing some of the advantages of recombinant antibodies. This has changed dramatically with the application of SpyTag/SpyCatcher protein ligation technology to recombinant antibodies. This biochemical format conversion enables the almost instantaneous generation of a versatile family of formats, including rabbit, mouse and human synthetic immunoglobulins, as well as site-specific labeling with horseradish peroxidase (HRP), biotin, and other tags^53^. In particular, a bivalent modular antibody carrying three HRP molecules has proven to be a much more sensitive, simpler and faster immunoblotting approach compared to standard protocols based on secondary antibodies^53^, significantly improving detection of ADPr^21,47^.

Here, we extend the scope of our serine ADPr technology to poly-ADPr, combining it with a proven phage display recombinant antibody platform and the SpyTag protein ligation system to develop sensitive and specific poly-ADPr antibodies. This advance enables versatile, precise and consistent detection across a variety of experimental applications, such as immunofluorescence, immunoblotting and immunoprecipitation. In this study, we present the characterization and validation of new poly-ADPr recombinant antibodies, reasoning that the modular nature of this antibody, as well as our previously introduced complementary antibodies, will enable more versatile and sensitive ADPr detection. Despite recent advances, many aspects of ADP-ribosylation remain poorly understood, with outstanding questions regarding its substrate specificity, spatial and temporal regulation. Furthermore, recent discoveries of novel ADP-ribosylation readers, writers, and erasers promise to reshape our understanding of this modification and its broader impact on cellular physiology.

## Material and Methods

### Resource availability

#### Lead contact

Further information and requests for resources and reagents should be directed to and will be fulfilled by the lead contact, Dr. Ivan Matić

#### Material availability

Antibodies, Reagents and BiSpyCatcher

The α-Poly-ADPr antibodies generated in this study (AbD64138 and AbD64235) are not yet commercially available but requests can be forwarded to the lead contact. Previously published antibodies converted to the SpyTag formats (AbD43647) and SpyCatchers are available through BioRad Laboratories (unconjugated, requires conjugation to a SpyCatcher: TZA020; HRP-conjugated, ready-to-use: TZA020P)

Uncoupled SpyTagged antibodies were conjugated as described below to following SpyCatchers: bivalent-HRP Format (BiSpyCatcher2 #TZC002P, Bio-Rad); bivalent synthetic IgG Format mouse (Mouse IgG2a-FcSpyCatcher3 # TZC012, Bio-Rad); bivalent synthetic IgG Format rabbit (Rabbit IgG-FcSpyCatcher3 # TZC013, Bio-Rad)

Commercial antibodies and reagents used in this study: α-Poly/Mono-ADP Ribose (D9P7Z) (Cell Signaling Technology, #89190), α-Poly (ADPr-ribose) monoclonal antibody (10H) (Enzo Life Sciences, ALX-804-220-R100), α-Poly-ADP-ribose binding reagent (Merck, MABE1031)

### Experimental Model

#### Cell Culture and drug treatments

U2OS cell lines were obtained, authenticated by STR profiling and confirmed mycoplasma free by ATCC cell line authentication service. Cells were cultured in Glutamax-DMEM supplemented with 10% bovine serum and 100U/ml penicillin/Streptomycin at 37°C and 5% CO_2_.

To induce DNA damage, the cell medium was aspirated and replaced with 37°C complete DMEM containing 2mM H_2_O_2_ for 10 min. For PARP inhibition (Olaparib, Cayman Chemical) cells were treated with 1 μM Olaparib in complete DMEM for 30 min prior to DNA damage treatment, followed by 10 min 2mM H_2_O_2_ & 1μM Olaparib treatment.

### Method details

#### Antibody SpyCatcher Coupling

All Bio-Rad antibodies have been obtained unconjugated in a monovalent format and were conjugated to BiSpyCatchers inhouse leading to bivalent HRP conjugated or bivalent synthetic IgG antibody formats. In brief antibody stocks were diluted to 1 mg/ml in PBS and incubated with 1/10th the volume of unconjugated antibody with BiSpyCatcher for 1 hr at RT, i.e. 10 μl of BiSpyCatcher was added to 100 μl of 1mg/ml unconjugated antibody. Afterwards they were aliquoted and stored at -20°C.

#### Expression and purification of recombinant proteins

Proteins were expressed in *Escherichia coli* (*E.coli*) and purified essentially as previously described^54^, but with addition of a size-exclusion chromatography purification using HiLoad 16/60 Superdex 75 column. PARP1 wild-type was expressed and purified as previously reported^54,55^. PARG was purified as described^56^. HPF1 was expressed and purified as reported^57^.

### In vitro Modification of Proteins

Recombinant PARP1 protein (0.5 μM) was incubated with 2 mM NAD^+^ in the presence of HPF1 (5 μM). The reaction buffer contained 50 mM Tris-HCl pH 8.0, 100 mM NaCl, 2 mM MgCl_2_ and 1 ng/μL sonicated DNA. Reactions were allowed to proceed for at 30 min at RT, before being stopped by the addition of 1 μM Olaparib. To create mono- and poly-ADP-ribosylated protein samples were split and one half was further incubated with 3 μM PARG for 1 hr at RT and stopped by the addition of Laemmli buffer. Samples were boiled and run on an 8% Bis-Tris gel (Invitrogen).

For Dot-blotting the same reaction as above was performed without splitting the reaction after stopping with Olaparib. No loading buffer was added. A dilution series of the sample was performed using reaction buffer and the same volumes (1 μl) of the probe was pipetted onto a dry nitrocellulose membrane. The membrane was left to dry and was proceeded with 5% non-fat dried milk blocking and regular immunoblotting steps as described below.

#### SDS Sample Preparation and Immunoblotting

U2OS cells were treated as indicated, lysed in SDS buffer (4% SDS; 50 mM HEPES pH 7.2, 150 mM NaCl, 5 mM MgCl_2_) and incubated for 5 min at RT with recombinant benzonase (smDNase, 750U per sample). Samples were spun at max. speed for 10 min and supernatant was transferred into fresh tube. Sample concentration was measured using a Pierce Dilution-Free BCA Protein Assay Kit (#A55864, ThermoScientific). Samples were prepared with NuPAGE LDS sample buffer with final concentration of 5 mM DTT (Sigma) and loaded on 4-12% Bis-Tris gel (Invitrogen). After SDS-Page the gels are transferred onto PVDF membranes (Amersham) using wet transfers at 90 mA overnight on ice. The precooled transfer buffer is 1x NuPAGE transfer buffer (Invitrogen), 20% ethanol in water.

For immunoblotting membranes were blocked in 5% non-fat dried milk before primary antibodies were added. Primary Antibodies (commercial antibodies and HRP-formats of Bio-Rad conjugated antibodies) were prepared in milk using concentrations found in figures. The membranes were incubated overnight at 4°C before being washed in TBS-T (25 mM Tris-HCl pH 7.5, 150 mM NaCl, 0.05% Tween-20) 3×10 min. Commercial antibodies (Enzo, Merck, Cell Signaling Technologies) were incubated with a secondary antibody (Anti-mouse IgG HRP-conjugated secondary Amersham Cat# NA931V, Anti-rabbit IgG HRP-conjugated secondary Merck Cat# GENA934-1ML) for 1 hr at RT, followed by 3×10 min TBS-T wash. All membranes were developed using ECL Select Western Blotting Detection Reagent (Amersham).

#### Immunofluorescence

U2OS cells were grown on sterile glass coverslips in 24-well plates. treated as indicated and fixed for 20 min with Methanol at -20°C. The fixed cells were washed twice with PBS, and blocked for 1hr, RT with 5% Normal-Goat-Serum in PBS supplemented with 0.3% Triton. Primary antibodies (commercial antibodies and synthetic IgG formats of Bio-Rad conjugated antibodies) were diluted as indicated and incubated overnight at 4°C in 1x PBS, 1% BSA and 0.3% Triton. Coverslips were washed in PBS 3×5 min before adding the secondary antibody and DAPI, again in 1x PBS, 1% BSA and 0.3% Triton for 1 hr, RT protected from light. Coverslide were washed again in PBS 3×5 min before mounted on microscopy slides using Prolong Diamond Antifade Mountant (ThermoFisher). Cells were scanned for immunofluorescence using a Leica SP8-X inverted laser-scanning confocal microscope.

## Results

### Generation of a serine poly-ADPr peptide

Previously, we established the PARP1/HPF1 complex^20,57,58^ as an efficient and specific biocatalytic tool for the rapid and scalable generation of mono-ADPr peptides, enabling the development, validation and characterization of site-specific, mono-ADPr and pan-ADPr antibodies^46,47^. Here, we aimed to adapt our strategy to generate modular antibodies specific for poly-ADPr. Inspired by the possibility of extending the initial ADP-ribose attached to serine of the peptide^40^, we employed a simple protocol that does not require separation of different ADP-ribosylated peptide species, reasoning that this approach would be sufficient for the development of poly-ADPr antibodies. Specifically, we generated and purified the mono-ADPr peptide from proteins using C18 StageTips^46^ and subsequently added PARP1 without HPF1 to extend the mono-ADPr modification to di- and tri-ADPr. This process proved to be as scalable as the previously described serine mono-ADPr peptide generation method^46^. The resulting peptides were then employed in a phage display screen, leading to the identification of several positive clones. From these, we selected the two most promising candidates for further validation and comparison, as detailed in the following sections (AbD64138, AbD64235).

### Modular Antibodies for versatile, specific and sensitive detection of Poly-ADPr

Given the demonstrated value of the SpyTag system in the generation of modular antibodies^46,47^, we also chose to implement it for our new set of poly-ADPr antibodies. As illustrated in the scheme (Fig. 1), the monovalent poly-ADPr antibody can be easily converted into a bivalent format in a fast highly reliable reaction. This conversion adds an avidity effect in addition to the high affinity of the antibody, further enhancing signal intensity and specificity. Depending on the application, a variety of catchers can be chosen for coupling. For example, an HRP-catcher can be used, resulting in an HRP-coupled bivalent poly-ADPr antibody for immunoblotting. This eliminates the need for a secondary antibody, reducing costs, antibody incubation time and increasing signal intensity, as demonstrated previously^47,53^. Alternatively, for applications not reliant on HRP, such as immunofluorescence, a catcher fused with the mouse or rabbit Fc region can be selected, creating a synthetic IgG antibody that, like a regular IgG, is detectable by fluorophore-coupled secondary antibodies.

**Figure 1:**
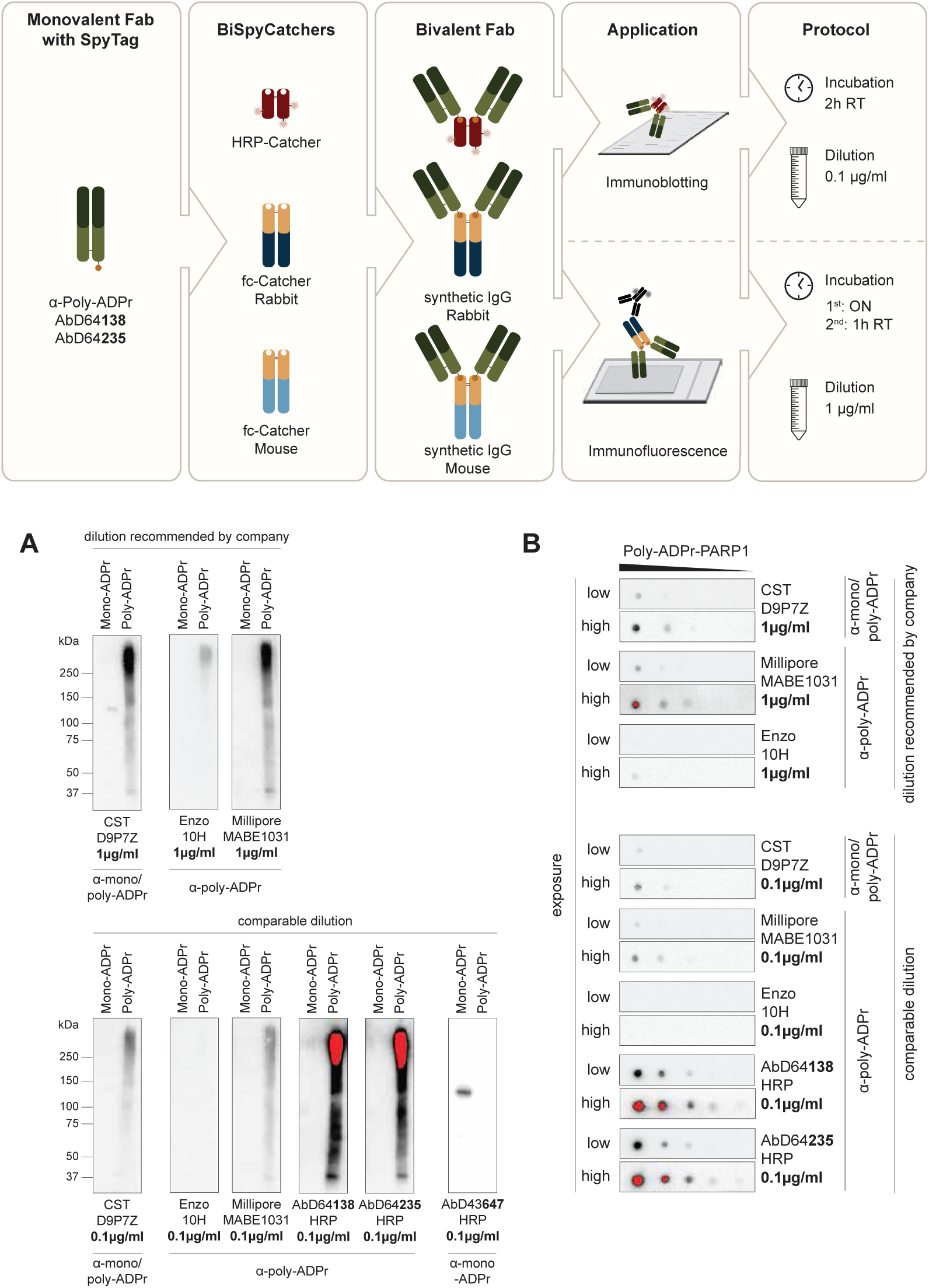
Modular Antibodies for versatile, specific and sensitive detection of Poly-ADPr. Scheme describing the coupling process of monovalent SpyTagged antibody, using various BiSpyCatchers to create a variety of formats for multiple approaches. **A:** *In Vitro* modified recombinant PARP1, treated w/o PARG to create Poly-/ and Mono-ADPr PARP1 immunoblotted with new anti-Poly-ADPr antibodies in comparison to other available antibodies and reagents. Red represents oversaturated signal. **B:** *In Vitro* modified recombinant PARP1 dot-blotted in dilution series and incubated with new anti-Poly-ADPr antibodies in comparison to competitors. Red represents oversaturated signal.

After coupling, we first assessed the specificity of the poly-ADPr antibodies (AbD64138, AbD64235) to ensure there was no off-target binding to other forms of ADPr. To test this, we took advantage of a key feature of serine ADPr: the inability of PARG to cleave the bond between serine and the initial ADP-ribose, despite its ability to hydrolyze tyrosine and aspartate/glutamate ADPr^21,59^. Specifically, we automodified PARP1 in the presence of HPF1, resulting in nearly complete serine Poly-ADPr of PARP1. Addition of PARG to a portion of the reaction completely reduced Poly-ADPr-PARP1 to mono-ADPr-PARP1^46^, resulting in pure mono- and poly-ADPr standards for antibody validation. As expected, our mono-ADPr antibody^47^ recognized exclusively mono-ADPr (Fig. 1A). Our newly developed poly-ADPr antibodies did not detect any mono-ADPr, but produced an intense signal for poly-ADPr PARP1. In contrast, pan-ADPr antibodies, such as D9P7Z from Cell Signaling Technology (CST), detected both mono- and poly-ADPr (Fig. 1,A). We also compared our antibodies with other commercially available poly-ADPr reagents, including 10H (Enzo) antibody and the WWE-based poly-ADPr reagent from Merck (MABE1031)^42^. All poly-ADPr probes were shown to be specific for poly-ADPr, although 10H only detected long ADPr polymers as previously reported^60^ (Fig. 1, A). Like MABE1031, our new antibodies were able to detect the full range of poly-ADPr chains in immunoblotting. To ensure a fair comparison of signal intensities, we tested the reagents at two different dilutions, starting with those recommended by the manufacturers/publications for optimal performance. However, we found that using our antibodies at the same concentration (1 µg/ml) led to signal bleaching before detection. Consequently, we lowered the concentration by a factor of ten, which allowed for signal detection without compromising the poly-ADPr detection by commercially available reagents. Despite this adjustment, the sensitivity of our antibodies is so high that oversaturation occurred after just one second of exposure.

To allow for a more accurate comparison without signal saturation, we performed a dot blot using a 1:2 dilution series of poly-ADPr PARP1 (Fig. 1,B). Again, using antibody dilutions as recommended by the manufacturer and a common dilution of 0.1 µg/ml, we compared the signal intensities of our new tools to those of other reagents. The results showed that our new antibodies provided significantly higher signal intensity compared to the MABE1031 reagent and D9P7Z antibody. The 10H antibody did not produce any detectable signal at this dilution.

These results demonstrate a high specificity for poly-ADPr of the new antibodies, with no detection of mono-ADPr, confirming that our antibodies selectively recognize poly-ADPr chains.

### Sensitive detection of Poly-ADPr targets in cells

After establishing their poly-ADPr specificity and sensitivity in *in vitro* biochemical reactions, we applied the new antibodies to cell lysates to evaluate their performance on cellular poly-ADPr targets and assess potential off-target binding. To induce high levels of poly-ADPr, cells were treated H_2_O_2_ for ten minutes and additionally treated with the PARP1 inhibitor Olaparib. We used optimal antibody dilutions for all probes to ensure peak performance. First, we demonstrated that there was no nonspecific binding to proteins, as shown in lanes 2 and 4 of each blot, where no signal was detected in olaparib-treated samples (Fig. 2,A). Second, even when using ten times higher amounts of commercially available reagents, our new poly-ADPr antibodies still exhibited superior signal intensity. Third, our antibodies also improved the detection of poly-ADPr in unperturbed cells, where its generation has been shown to be induced by unligated Okazaki fragments during DNA replication^61^, even with low exposures, as seen in lane 1.

**Figure 2:**
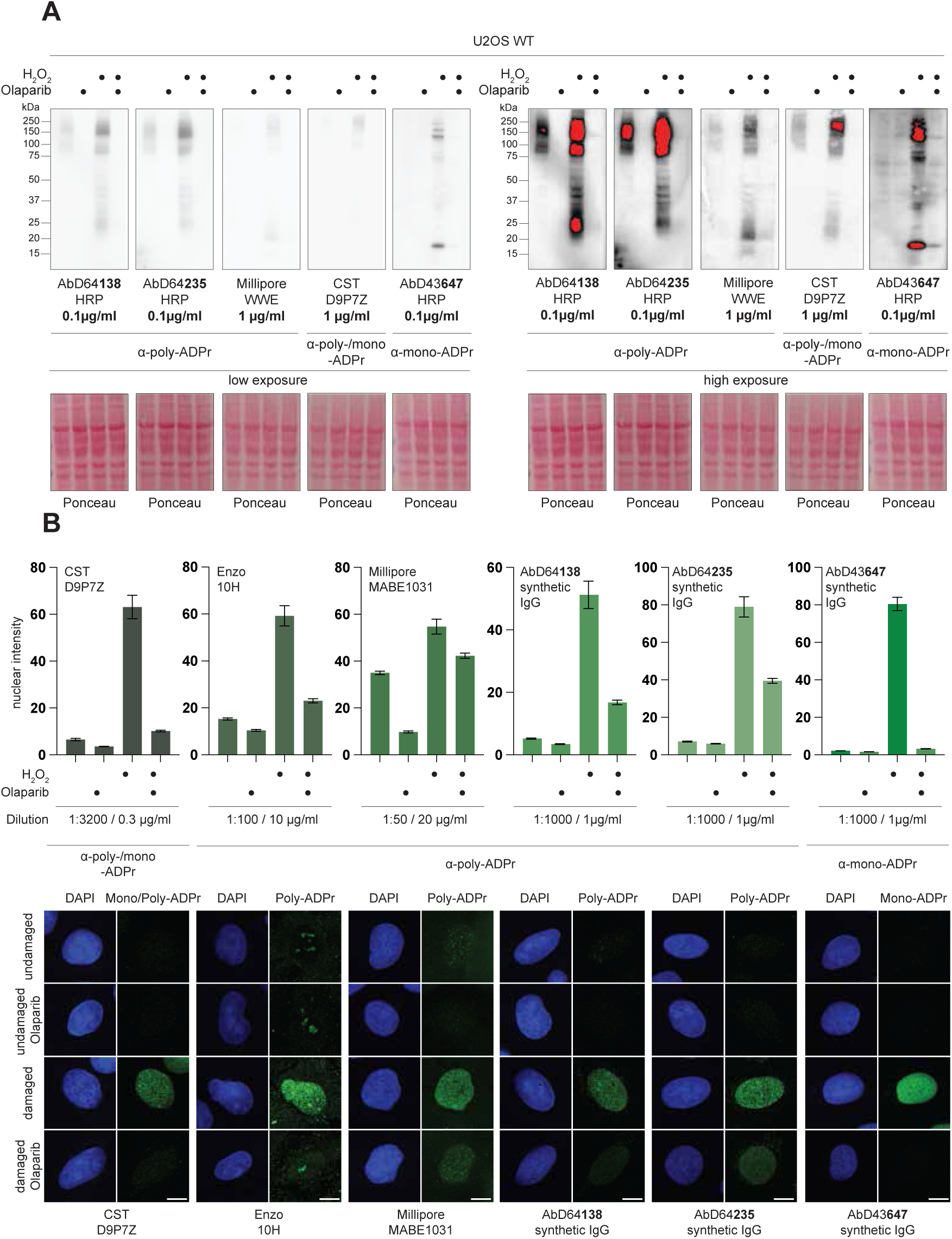
New highly sensitive Poly-ADPr-specific Antibodies. **A:** Immunoblotting of U2OS cells treated or not with H_2_O_2_, w/o Olaparib, lysed in SDS buffer and proceeded with immunoblotting. Antibodies and Reagents used in indicated concentration recommended by manufacturer or publication. HRP-format was used for new anti-poly-ADPr antibodies. Red represents oversaturated signal **B:** Immunofluorescence images of U2OS cells treated or not with H_2_O_2_, w/o Olaparib followed by methanol fixation. Antibodies and Reagents used in indicated concentration recommended by manufacturer or publication. Synthetic IgG format was used for new anti-poly-ADPr antibodies. Error bar represents SEM. Scale bar represents 10 μm.

We then tested the specificity of our antibodies in cells using immunofluorescence and the synthetic IgG format. Again, we applied optimal antibody dilutions and tailored microscope settings for each reagent to achieve high performance. While intensity levels are not directly comparable, the focus was on the specificity of the signal and the signal-to-noise ratios. Our poly-ADPr antibodies AbD64138 and AbD64235 provided a superior signal-to-noise ratio in untreated versus H_2_O_2_-treated cells (Fig. 2,B), outperforming other poly-ADPr reagents. The signal specificity was comparable to the poly-/mono-ADPr antibody (CST) and our mono-ADPr reagent (AbD43647), with antibody AbD64138 performing slightly better over AbD64235.

In conclusion, these experiments validate the high signal-to-noise ratio of our new tools for both immunoblotting and immunofluorescence, demonstrating their versatility and potential for multiple applications.

### SpyTag System Enables Flexible Co-Detection of Mono- and Poly-ADPr In Cells

Finally, to assess the potential for simultaneous use of our poly-ADPr antibodies alongside our previously published mono-ADPr antibodies, we performed co-staining of both poly-ADPr (AbD64138) and mono-ADPr (AbD43647) in cells (Fig. 3,B). For this, we coupled the poly-ADPr antibody with a mouse Fc catcher and the mono-ADPr antibody with a rabbit Fc catcher. To prevent channel bleed-through, we used two fluorophores with distinct emission peaks and employed line sequencing during imaging. The results demonstrate that, following a ten-minute H_2_O_2_ treatment, both mono- and poly-ADPr can be detected simultaneously in the same cells. Notably, the staining patterns revealed distinct distributions: the mono-ADPr signal appeared more homogenously distributed throughout the nucleus, while the poly-ADPr signal exhibited a more punctate pattern, forming small foci.

**Figure 3:**
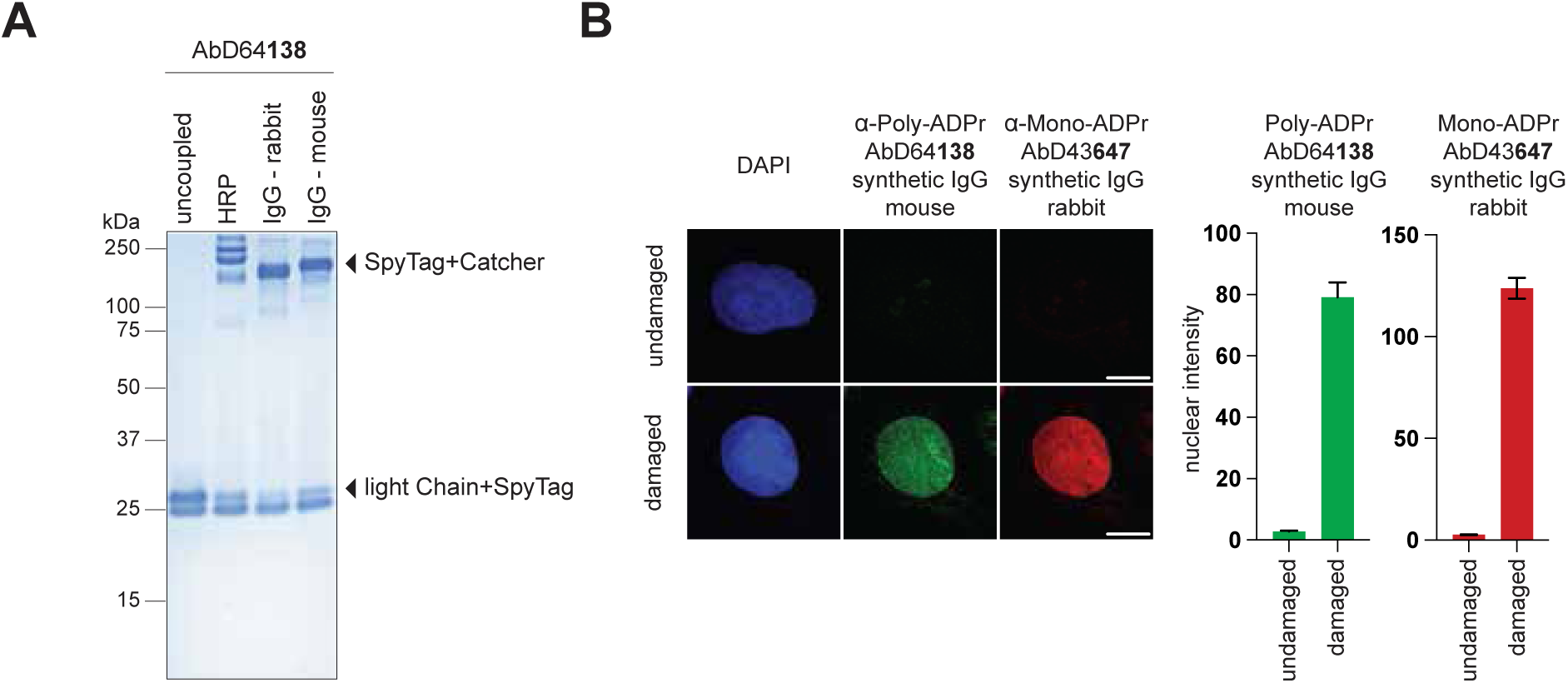
Versatile SpyTag System Enables Flexible Co-Detection of Mono- and Poly-ADPr In Cells. **A:** Coomassie Staining of the same AbD64138 antibody in monovalent unconjugated format (first lane) and bivalent conjugates formats (lane 2-4). **B:** Immunofluorescence co-staining of mono- and poly-ADPr in the same cell using synthetic mouse/rabbit IgG antibody formats.

The ability to co-stain both forms of ADPr enhances the depth of analysis in immunofluorescence, allowing for the use of smaller sample sizes and reduced experimental times. Additionally, the flexible coupling of different IgG species to either poly- or mono-ADPr antibodies provides (Fig.3, B) versatility for co-staining with other targets, such as full-length PARP1, facilitating broader applications in cellular analysis.

To summarize the results of our antibody validation, we demonstrate superior performance in both signal intensity and specificity for immunoblotting and immunofluorescence applications. While both of our antibodies are highly specific for poly-ADPr and exceptionally sensitive, especially in immunoblotting, AbD64138 shows slightly better performance compared to AbD64235.

## Discussion

Despite decades of research on ADPr and recent technological advances^29^, the availability of specific and sensitive tools for studying all forms of this challenging PTM remains limited. This study aimed to extend the reach of our serine ADPr technology, which enabled the development of the first site-specific ADPr antibodies as well as specific and sensitive mono-ADPr antibodies^21,46–49^, to poly-ADPr, resulting in a highly sensitive, specific and versatile modular antibody. As a technology, Serine ADPr offers several advantages inherent to the nature of the physiological reaction. First, PARP1 is the most catalytically active PARP enzyme, strongly activated by DNA breaks both in cells and in biochemical reactions. Therefore, the modification is highly scalable, allowing reactions to be upscaled to milligram quantities suitable for antibody panning. Second, HPF1 enables not only precise targeting of PARP1 activity toward serine residues, but also modification of peptides rather than just PARP1 automodification^20^. Third, the biological interplay between serine ADPr and canonical histone marks^62^ can be harnessed to program the HPF1/PARP1 complex, directing it to target a specific serine using phosphorylation as an unconventional protecting group^46^. Fourth, a key advantage of serine-ADPr is its chemical stability: the O-glycosidic linkage remains intact across a broad range of pH conditions and withstands heat exposure up to 95°C. This stability facilitates efficient handling and minimizes sample loss during experimental procedures, a problem observed with other forms of ADPr, particularly aspartate/glutamate ADPr^21,63^. Fifth, while the poly-ADPr eraser PARG can cleave the bonds between ADP-ribose and tyrosine, aspartate and glutamate^21,59^, it does not remove serine ADPr. Here, beyond peptide generation for phage display and antibody screening, we harness the resistance of serine ADPr to PARG to achieve nearly complete modification of PARP1 at the protein level, either by poly-ADPr or, upon PARG treatment, of mono-ADPr. This enables comprehensive antibody specificity testing, ensuring the selection of clones with high specificity for poly-ADPr and minimal off-target binding.

The newly developed poly-ADPr antibodies presented in this study offer several key advantages. They enable the detection of a broad range of poly-ADPr, from short to long polymeric chains, as demonstrated by full signal coverage in immunoblotting analysis. The modular nature of these antibodies, derived from the SpyTag technology results in strong signal intensity, especially in immunoblotting, and offers versatility across multiple applications, including immunofluorescence and immunoprecipitation, allowing for the detection of poly-ADPr even under untreated conditions. The modular SpyTag system, as illustrated previously for our mono-ADPr and site-specific antibodies^47^, provides significant benefits. The reaction used to couple various catchers to the monovalent antibodies is fast, efficient, and highly reliable. For immunoblotting, the HRP Catcher eliminates the need for a secondary antibody, significantly increasing signal intensity, reducing costs by lowering the required antibody amount per experiment and shortening the overall immunoblotting procedure. Additionally, the SpyTag system enables the creation of our poly-ADPr antibodies as synthetic immunoglobulins, which can be used in immunofluorescence assays. This flexibility allows for the selection of species-specific antibodies (e.g., mouse, rabbit, or human) tailored to experimental needs, allowing co-staining with other ADPr- or protein-specific antibodies.

Beyond their high affinity and modularity, the antibodies described in this study also offer high reliability. As monoclonal recombinant antibodies, they ensure consistent quality and reproducibility, unlike previously available polyclonal poly-ADPr antibodies, which were subject to batch variability and eventual discontinuation. This homogeneity and consistency further reinforce their utility as a long-term tool for ADPr research.

The most critical factor when selecting a tool to study ADPr is that its effectiveness is inherently dependent on the research question being asked. Choosing the appropriate tool is essential to obtaining meaningful and accurate insights. Given the high variability of ADPr arising from modifications at different sites, in diverse forms, by various proteins across multiple cellular signaling pathways, multiple complementary tools are required to study this PTM in an in-depth manner. Broad yet ADPr-specific tools, such as the highly sensitive poly-/mono-ADPr antibody from Cell Signaling Technology, allow for the general detection of ADPr. However, their inability to distinguish between different ADPr forms can complicate result interpretation, especially as it is emerging that mono- and poly-ADPr act as distinct signals, even within pathways where they co-occur^21,47^. In this study, we aimed to develop antibodies that provide both broad detection and high specificity, fully complementing our previously developed mono-ADPr antibodies^46,47^ to enable a clear differentiation between mono- and poly-ADPr. The tools presented here contribute to expanding the available toolkit for ADPr research at this level. To further investigate mono- and poly-ADPr in greater depth, additional specialized tools may be necessary. For instance, site-specific mono-ADPr and poly-ADPr antibodies may enable a more detailed examination of ADPr modifications, while engineered protein domains such as WWE fused to an Fc chain^39,43,64^, offer an alternative approach to studying certain forms of poly-ADPr.

In conclusion, we present the first application of our technology for generating poly-ADPr-specific antibodies. As with the development of site-specific and mono-ADPr antibodies, we incorporated serine ADPr at nearly every phase of the antibody generation process, from precisely modified peptide antigens to multiple validation steps and in-depth characterization. Obtaining purely mono- and poly-ADP-ribosylated PARP1 was particularly useful for unambiguously establishing the specificity of the new antibodies. In this process, phage display and the SpyTag protein ligation system were essential for generating numerous initial candidate clones and for conferring modularity to the final, validated antibodies, increasing their versatility and sensitivity. We anticipate that these new tools, along with previously developed reagents, will play key roles in deciphering the distinct roles of the two primary forms of ADPr - its monomeric and polymeric signals - in various signaling pathways.

## Acknowledgements

This work was funded by the Max Planck Society, the Deutsche Forschungsgemeinschaft (DFG, German Research Foundation) under Germanýs Excellence Strategy (CECAD, EXC 2030-390661388) and by the European Research Council (ERC-CoG-864117) to I.M.

## Author Contributions

I.M. conceived the project. H.D. and I.M. designed research and wrote the manuscript. H.D. performed all the experiments.

